# Microbiome Features Associated with Performance Measures in Non-Athletic and Athletic Individuals

**DOI:** 10.1101/2023.06.10.544446

**Authors:** Kinga Humińska-Lisowska, Kinga Zielińska, Jan Mieszkowski, Monika Michałowska- Sawczyn, Paweł Cięszczyk, Paweł P Łabaj, Bartosz Wasąg, Barbara Frączek, Anna Grzywacz, Andrzej Kochanowicz, Tomasz Kosciolek

## Abstract

The influence of human gut microbiota on health and disease is now well established. Therefore, it is not surprising that microbiome research has found applications within the sports community, hoping to improve health and optimize performance. Every comparison study found new species or pathways that were more enriched in elites than in sedentary controls. In addition, sport-specific and performance-level-specific microbiome features have been identified. However, the results remain inconclusive and indicate the need for further assessment. In our study, we conducted two interventions, anaerobic (Wingate Anaerobic Tests (WAnTs)) and aerobic (Bruce Treadmill Test), on fit non-athletic but physically active controls and athletic strength (training professional sports which increase anaerobic efforts such as weightlifting, powerlifting, and bodybuilding) and endurance (training professional sports which increase aerobic efforts such as race walking, long-distance running, or ski running) individuals to compare sport-specific and training-specific responses. While we did not identify any differences in alpha and beta diversity or significant differential abundance of microbiome components at baseline, we noted that one-third of the species identified were unique to either group. Longitudinal analysis of samples (pre- and post-intervention) revealed an abundance of *Alistipes communis* in the strength group during the WAnT intervention and 88 species with notable differences between groups during the Bruce Test intervention. SparCC recognized *Bifidobacterium longum* and *Bifidobacterium adolescentis*, short-chain fatty acid producers with probiotic properties, as species strongly associated with VO_2_max. Ultimately, we identified several taxa with different baseline abundances and longitudinal changes when comparing individuals based on their VO_2_max, average power, and maximal power parameters. Our results confirmed the health status of the individuals involved in the study, based on previous assumptions about microbiome health. Furthermore, our findings indicate that microbiome features are associated with better performance measures previously identified in elite athletes.

## 1. Introduction

The human microbiota refers to 10-100 trillion symbiotic microbial cells, mainly human gut bacteria, and the set of genes they possess is termed the human microbiome [1]. Thanks to initiatives such as the Human Microbiome Project [2] or the American Gut Project [3], there has been an increasing interest in the functions of the microbiome and its role in maintaining human health [4]. The human microbiome is influenced by a wide range of host factors, such as sleep [5], diet [6], and age [7], but also by many other factors (host genome, stress, illness, and drugs) [8]. Any disruption to the microbiome homeostatic state, collectively known as dysbiosis, is thought to have a number of consequences and may be one of the factors linked to the diseas [9].

However, not all changes in the microbiome composition are always negative. Some indicate the human body’s response to external stimuli, and its character is aimed at adapting to the strength of the acting factor. One factor affecting human body function that may cause microbiome changes is physical activity. The beneficial effects of physical activity are widely known, well established, and examined. Exercise may induce both acute and chronic alterations in the microbiome, and is widely agreed to be a positive and vital stimulus affecting health [10]. Alterations in the health-associated microbiome are also known. Research has shown that microbiome activity may be linked to sports performance [11], and could eventually guide the optimization of each athlete’s abilities by changing microbiome composition for targeted performance improvement. Apart from sports, this could have practical implications in treating health disorders, such as obesity and cardiovascular diseases [12].

Some studies have suggested that athletic gut microbiota has higher levels of alpha diversity and health-supporting gut bacteria [13]. While studies remain inconsistent, species such as *Faecalibacterium prausnitzii, Roseburia hominis, Akkermansia muciniphila* and *Eubacterium rectale* are generally cited as having a positive influence on health and performance [14]. Athletes also tend to have an enrichment of short-chain fatty acids (SCFA) modulating substrate metabolism at various organ sites [15]. Bacterial species that influence SCFA production or breakdown, such as *Veillonella atypica*, have been shown to positively influence marathon running time [16]. Furthermore, athletes are known to have several enriched pathways compared with sedentary individuals. These include amino acid and antibiotic biosynthesis and carbohydrate metabolism [17]. The features of a healthy athletic microbiome are not universal but are highly dependent on individuals and their sport characteristics. Endurance exercises are mainly associated with an aerobic type of physical activity and aerobic metabolic changes such as increased production of SCFAs and an abundance of inflammation-reducing microbiota [10]. In contrast, enhanced use of protein in strength sports decreased the occurrence of SCFA-producing bacteria [18].

Most studies are based on differences between elite athletes (healthy or not) and sedentary, often unhealthy, individuals. Because of that, it is difficult to identify which microbiome features are linked to health or strictly to sports performance. Furthermore, the results of current cross-sports studies are usually inconclusive [19] and in comparison to different controls, unlike athletic microbiome markers are determined. Our suggestion is that prolonged professional training in endurance or strength sports leads to physiological changes that may affect intestinal microbiome populations. Furthermore, we hint that long-term training will reveal diverse post-exercise microbiome responses, primarily dependent on the type of discipline practiced: endurance (training professional sports which increase aerobic efforts such as race walking, long-distance running, or ski running) or strength (training professional sports which increase anaerobic efforts such as weightlifting, powerlifting, and bodybuilding). In order to focus strictly on the performance-associated microbiome characteristics, we included only healthy individuals in our study. Specifically, we compared two sport-specific populations: training strength-associated sports (strength) and endurance sport disciplines (endurance) to physically active, but not professionally trained individuals. We conducted two interventions with maximal intensity: anaerobic (Wingate Anaerobic Tests (WAnTs)) and aerobic (Bruce Treadmill Test) to gain insights into gut microbiota alterations related to different exercise modalities. As a result, we were able to eliminate the “health factor” and focus solely on pre-exercise and post-exercise microbiome alterations linked to two different exercise modalities (anaerobic and aerobic), specific to each group (strength, endurance and control).

## 2. Materials and Methods

### 2.1. Experimental overview

This is a case-control study which consisted of two parts. The first was the measurement of anaerobic (Wingate Anaerobic Tests (WAnTs)) and aerobic (Bruce Treadmill Test) components of fitness, and the second was the assessment of fecal microbiome DNA composition. Each part of this study is described in detail below. A week before any testing, all participants attended a familiarization session to ensure that they were familiar with the testing equipment and study procedures. Healthy, physically active male volunteers participated in this study.

Based on purposive selection, the study participants were assigned to three groups: endurance athletes (n= 15), strength athletes (n=16), and the control group (n=21). Endurance athletes included in the study exhibited a minimum of 5 years of training, focusing on disciplines such as race walking, long-distance running (marathon, half marathon, ultrarunning), or ski running. Conversely, strength athletes included in the study demonstrated a minimum of 5 years of training in weightlifting, powerlifting, and bodybuilding. For the control population, only physically active men who did not participate in any organized training but were physically active (participating in amateur sports activities mainly in team sports, in the gym, or swimming, but only occasionally) were included. Exclusion and inclusion criteria are presented in Table 1. The subjects provided written informed consent before enrolling in the study. Before participation, the subjects were informed about the study procedures; however, they were not aware of the rationale and study aim, and therefore, were naive about the main scope of the study. To determine the appropriate sample size, a power analysis of the interactions between effects was performed using GPower ver. 3.1.9.2. The minimal total sample size for a medium effect size at a power of 0.8 and a significance level of 0.05, was calculated as 15 for each group. A professional physician examined all the volunteers before and after each test. A 14-day break was used between the anaerobic and aerobic testing.

**Table 1.**
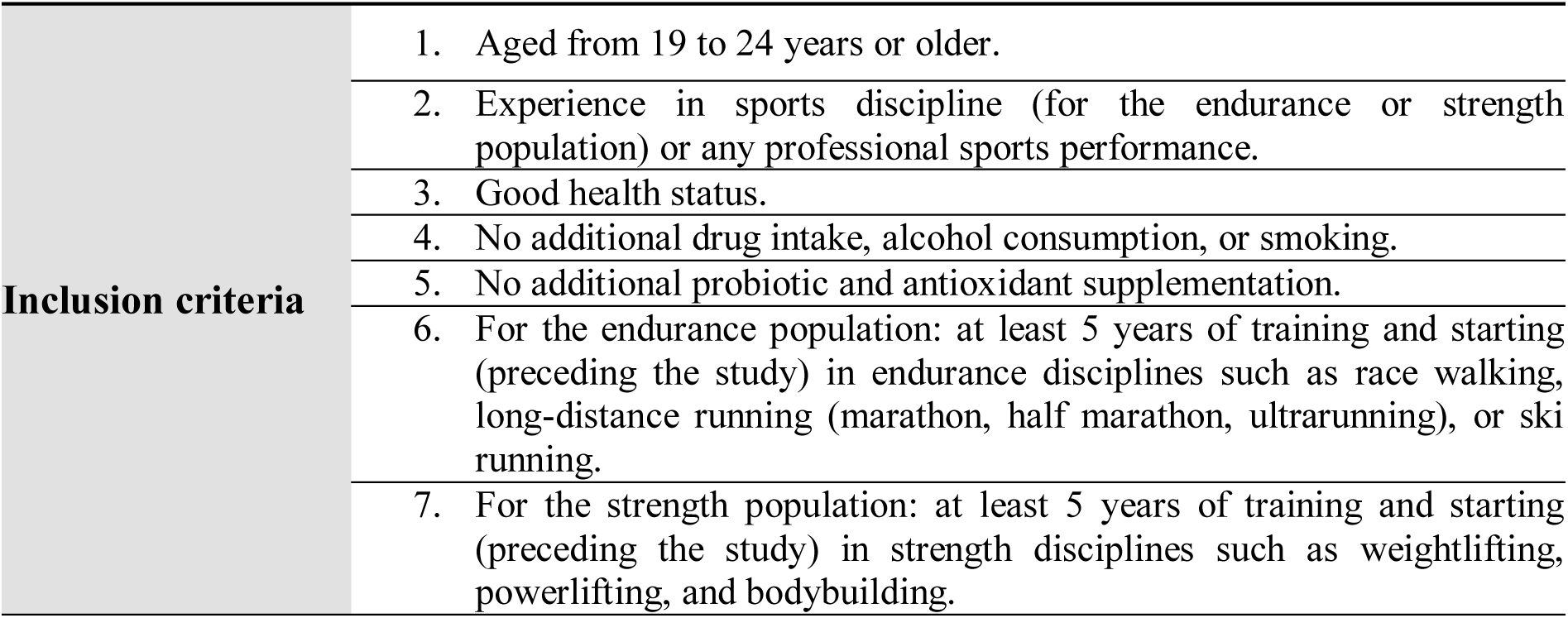
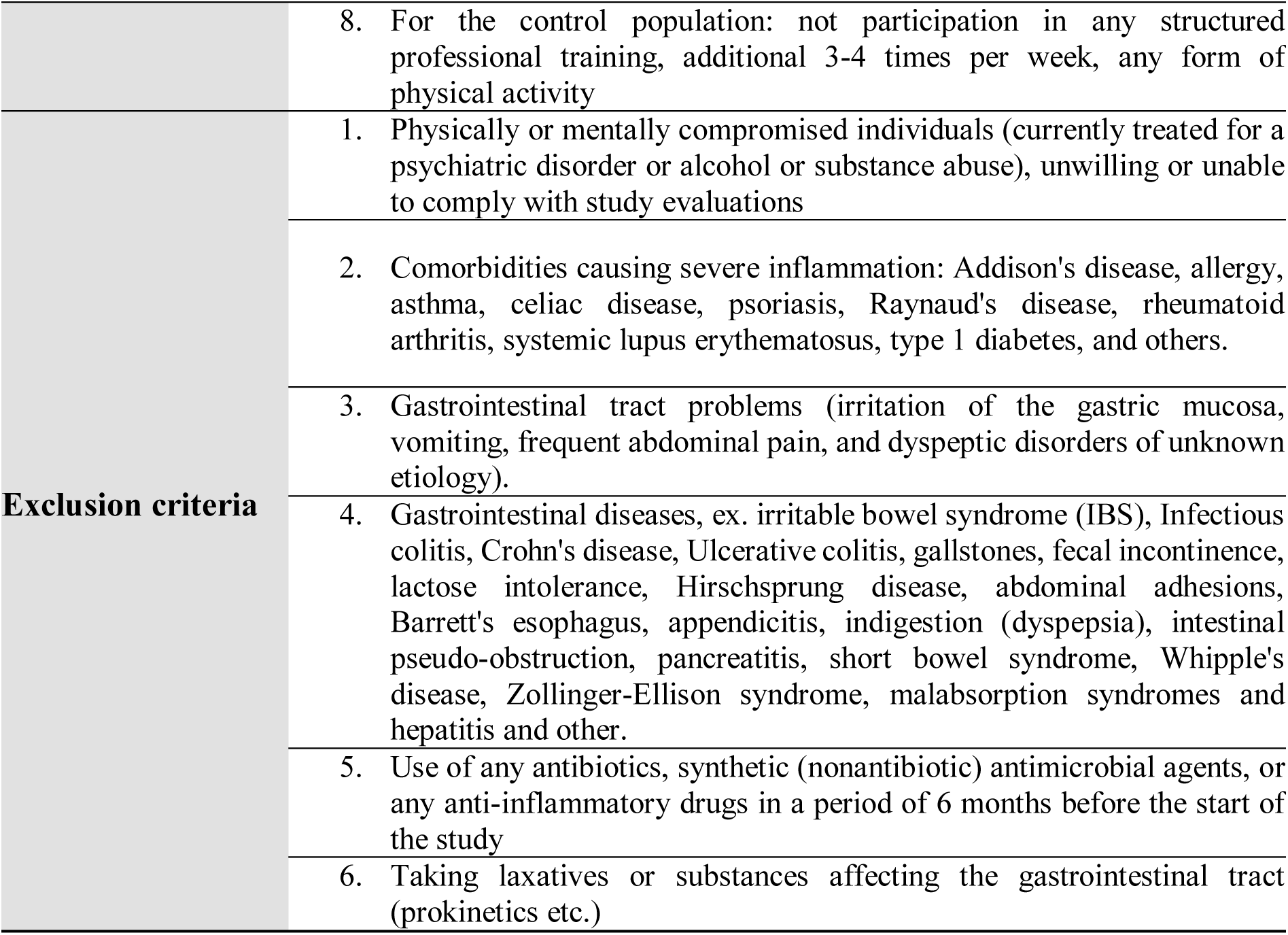
Eligibility criteria.

|  |  |
| --- | --- |
| **Inclusion criteria** | 1. Aged from 19 to 24 years or older. |
|  | 2. Experience in sports discipline (for the endurance or strength population) or any professional sports performance. |
|  | 3. Good health status. |
|  | 4. No additional drug intake, alcohol consumption, or smoking. |
|  | 5. No additional probiotic and antioxidant supplementation. |
|  | 6. For the endurance population: at least 5 years of training and starting (preceding the study) in endurance disciplines such as race walking, long-distance running (marathon, half marathon, ultrarunning), or ski running. |
|  | 7. For the strength population: at least 5 years of training and starting (preceding the study) in strength disciplines such as weightlifting, powerlifting, and bodybuilding. |
|  | 8. For the control population: not participation in any structured professional training, additional 3-4 times per week, any form of physical activity |
| **Exclusion criteria** | 1. Physically or mentally compromised individuals (currently treated for a psychiatric disorder or alcohol or substance abuse), unwilling or unable to comply with study evaluations |
|  | 2. Comorbidities causing severe inflammation: Addison's disease, allergy, asthma, celiac disease, psoriasis, Raynaud's disease, rheumatoid arthritis, systemic lupus erythematosus, type 1 diabetes, and others. |
|  | 3. Gastrointestinal tract problems (irritation of the gastric mucosa, vomiting, frequent abdominal pain, and dyspeptic disorders of unknown etiology). |
|  | 4. Gastrointestinal diseases, ex. irritable bowel syndrome (IBS), Infectious colitis, Crohn's disease, Ulcerative colitis, gallstones, fecal incontinence, lactose intolerance, Hirschsprung disease, abdominal adhesions, Barrett's esophagus, appendicitis, indigestion (dyspepsia), intestinal pseudo-obstruction, pancreatitis, short bowel syndrome, Whipple's disease, Zollinger-Ellison syndrome, malabsorption syndromes and hepatitis and other. |
|  | 5. Use of any antibiotics, synthetic (nonantibiotic) antimicrobial agents, or any anti-inflammatory drugs in a period of 6 months before the start of the study |
|  | 6. Taking laxatives or substances affecting the gastrointestinal tract (prokinetics etc.) |

All fitness analysis (Wingate Anaerobic Test and Bruce Trademill Test) were performed at Gdansk University of Physical Education (Gdansk, Poland) in October 2018. Before qualifying for the study, all participants completed a questionnaire on dietary preferences and physical activity to determine the population they would match (endurance, strength, or control population). Data on age, body composition (InBody 760 bioelectrical impedance analyzer), and height (anthropometer) of the individuals were collected during the initial visit.

All participants were instructed to maintain their everyday diet and were asked to refrain from vigorous exercise and avoid caffeine, alcohol consumption, and tobacco smoking during the week preceding the testing date, up to the last sample collection.

Food was not consumed during testing and water was available *ad libitum*. At the recruitment stage, the diet of the study participants was assessed. The results of the nutritional analysis, regarding the frequency of consumption of food products and food and the assessment of eating habits, were used to identify nutritional irregularities and to develop individual suggestions for modifying the diet in a rational direction, which was provided to all participants. In particular, nutritional recommendations were presented regarding the frequency of intake of food groups, the correct selection of food products (recommended products) and the principles of proper composition and size of meals and hydration. In addition, in order to ensure appropriate nutritional conditions, each participant received sample menus of a rational diet based on their nutritional preferences and in accordance with individual nutritional standards. Each study participant was given dietary recommendations for at least two weeks before the start of each exercise test until the last stool collection. In addition, on the day before each exercise test and on the day of the test, participants received meals at Gdansk University of Physical Education, established by the dietitian according to the individual energy expenditure of each participant.

None of the participants had any history of known diseases or reported medication intake (antibiotics or probiotics) due to illness six months before the experiment. Furthermore, none of the participants had direct contact with people with advanced bacterial, viral, or fungal diseases.

The study protocol was approved by the Bioethics Committee for Clinical Research of the Regional Medical Society in Gdansk (KB-27/18) and was conducted in accordance with the Declaration of Helsinki. All participants were informed of the possibility of withdrawing from the study at any time and for any reason, without any consequences.

All personal data were encoded and only information regarding the sports disciplines was accessible. The data were encoded starting from the recruitment stage to ensure participant anonymity and privacy.

During the examination, five individuals were excluded because they did not meet the study’s demands due to illness or injury.

#### Measurement of anaerobic components of fitness: double repeated lower body Wingate Tests

The double-repeated lower body WAnT was conducted on a cycle ergometer (Monark 894E, Peak Bike from Sweden) at the University of Physical Education and Sport, Gdansk Laboratory in October 2018. Every time the saddle height was adjusted, the final knee angle was approximately 170°–175°. The participants’ feet were held firmly in place (clips) to ensure complete contact with the pedals. Before any experimental testing, each individual completed a standardized warm-up on a cycle ergometer (5 min at 60 rpm, 1W/kg). Each participant was required to pedal with maximum effort for 30 s against a fixed resistive load of 75 g/kg of total body mass, as recommended by Bar-Or (1987) [20].

Before the test, the participants completed a warm-up that involved 5 min of arm cranking using a power output of 1 W/kg and crank rate of 60 rev/min. After the first test, participants had a 30 s break, and the WAnT was repeated in the same manner for the second time with maximum verbal encouragement from the beginning of each test [20].

During the WAnT test, certain variables were evaluated: peak power (W) and relative peak power (W/kg) were calculated as the highest single point of power output (recorded at 0.2 s intervals); mean power (W) and relative mean power (W/kg) were the average power output during the 30 s test.

#### Measurement of aerobic components of fitness: Bruce Treadmill Test

Each participant performed the Bruce Trademill Test two weeks after the WAnT (October 2018). The Bruce Protocol was performed on an electric treadmill (H/P/Cosmos, Germany). After standardizing the warm-up (5 min with 60% HR max), each participant performed running with increasing loads, including velocity and treadmill inclination. During the test, participants wore face masks connected to a pulmonary gas exchange analyzer (Quark CPET, Cosmed, Italy) and maximal oxygen uptake was measured. During the testing, participants performed specific test stages from 1 to 10 with increasing speed and velocity from a 10% Incline at 2.7 km/h (level 1) to a 28% Incline at 12.07 km/h (level 10). The test was stopped when the subject could not continue owing to fatigue or other conditions [21].

### 2.2. Samples Collection

The subjects’ performance was assessed using the WAnT (for maximal anaerobic exercise) and two weeks later, Bruce Treadmill Test (for maximal aerobic exercise) according to Bar-Or [19]. All participants collected fresh stool samples before and after each type of exercise at three time points: T0 (W0 or B0), in the morning fasting before the exercise test; T1 (W1 or B1), on the same day after the exercise test; and T2 (W2 or B2), the next morning on an empty stomach. The stool samples were collected in stool containers and immediately placed in thermal bags with cooling inserts and in a fridge until delivery to the laboratory within a maximum of a few hours, and immediately snap-frozen, and stored (−80 °C).

### 2.3. Samples Preparation

#### DNA isolation, quantitation and quantification

The frozen stool samples were thawed on ice. DNA was extracted from 200 mg of stool sample using a modified NucleoSpin® DNA Stool kit (Macherey-Nagel, Germany) via physical, mechanical, and chemical lysis (in March - May 2020). Mechanical lysis using a Bead Ruptor Elite (Omni International, US) was used to improve lysis. The following homogenization protocol was used:15sec homogenization with 6 ms speed × 5 times with 2 min of break after each 15 s. The concentration and purity of the total DNA isolates in the samples were measured spectrophotometrically using a NanoDrop (Thermo Scientific, USA) at wavelengths of A260 and A280. The degree of DNA degradation and potential contamination were analyzed using 1% agarose gels. Good quality and quantity DNA samples were snap-frozen, and stored (−80 °C). Prior to library preparation, the quantity of DNA was evaluated using a Qubit fluorometer and Qubit DNA BR Assay Kit in a Qubit® 3.0 Fluorometer (Thermo Fisher Scientific, USA). The final DNA concentration was standardized directly prior to library preparation.

#### Library preparation and sequencing

The library preparation procedure (performed in March – May 2022) followed the KAPA HyperPlus protocol (Roche, Basel, Switzerland). Stool-extracted DNA (100 ng of stool-extracted DNA) was used for the library preparation. Eighteen minutes of fragmentation at 37^0^C was applied to the library’s 300– 500 bp average fragment size. Indexed adapter ligation was performed using KAPA Universal dual-indexed adapters (Roche) at a concentration of 15 µM. To increase efficiency, adapter ligation was performed at 20^0^C for 1h. After ligation, the libraries were purified using a 0.8X KAPA Pure Bead (Roche, Switzerland) cleanup to remove unincorporated adapters. Six PCR cycles were performed during library amplification. The following cycling conditions were applied: 98 °C for 45 s; six cycles of 98 °C for 15 s, 60 °C for 30 s, and 72 °C for 30 s; final elongation at 72 °C for 1 min. Amplified fragments with adapters and tags were purified using KAPA Pure beads (Roche, Switzerland) at a 1:1 concentration.

Purified libraries were stored for up to 2 weeks at − 20 °C until sequencing. The quality of the libraries and fragment distribution were analyzed using a 4150 TapeStation System and D1000 reagents (Agilent Technologies, USA). Quality control of the purified libraries was performed for up to two weeks before the sequencing run.

Prior to sequencing in June – August 2022, all libraries were thawed on ice and normalized using the Qubit DNA HS Assay Kit (Thermo Fisher Scientific, USA) to a final concentration of 20 nM and mixed in equimolar concentrations to ensure an even representation of reads per sample. The libraries were sequenced on an Illumina NovaSeq sequencer (Novogene, Cambridge, MA, USA), and 150 bp paired-end reads were generated. Approximately 22 million 150 bp paired-end reads were generated per library.

### 2.4. Bioinformatics Analysis

Raw sequencing reads were preprocessed using Trim Galore! 0.6.10 [22]. Taxonomic profiles were calculated using Metaphlan4 [23], while functional and gene profiles were obtained using Humann 3.6 [24]. Between-group comparisons at all time points were performed using Qiime2 [25]: alpha and beta diversity, ANCOM, and LEfSe [26]. Longitudinal analysis was performed using the Qiime2 longitudinal plugin. Due to a limited number of samples collected by study participants prior to the WAnT test (n = 6) and a two-week break between the WAnT and Bruce tests, a comparative analysis was conducted between W0 and B0. Our reasoning presented in the Methods section showed that it was plausible to use the B0 samples instead of W0, as they were not statistically distant from one another, and not further than the post-intervention samples. We validated this approach by showing that, according to beta diversity, the few baseline WAnT samples were not further from the non-baseline WAnT or Bruce samples than the Bruce control samples. Correlations between microbiome features and metadata were calculated using MaAsLin [27]. The results were visualized using custom Python scripts. Authors who performed data analysis also had no access to information that could identify individual participants during or after data collection.

## 3. Results

### 3.1. Comparison of groups at baseline

The analysis of samples at baseline revealed no significant differences in alpha and beta diversity between the control, strength and endurance groups (Figure 1). Detailed alpha diversity (Shannon entropy, Simpson and Observed features) and beta diversity (Bray-Curtis and Jaccard) statistics are shown in Supplementary Tables 1 and 2.

**Figure 1.**
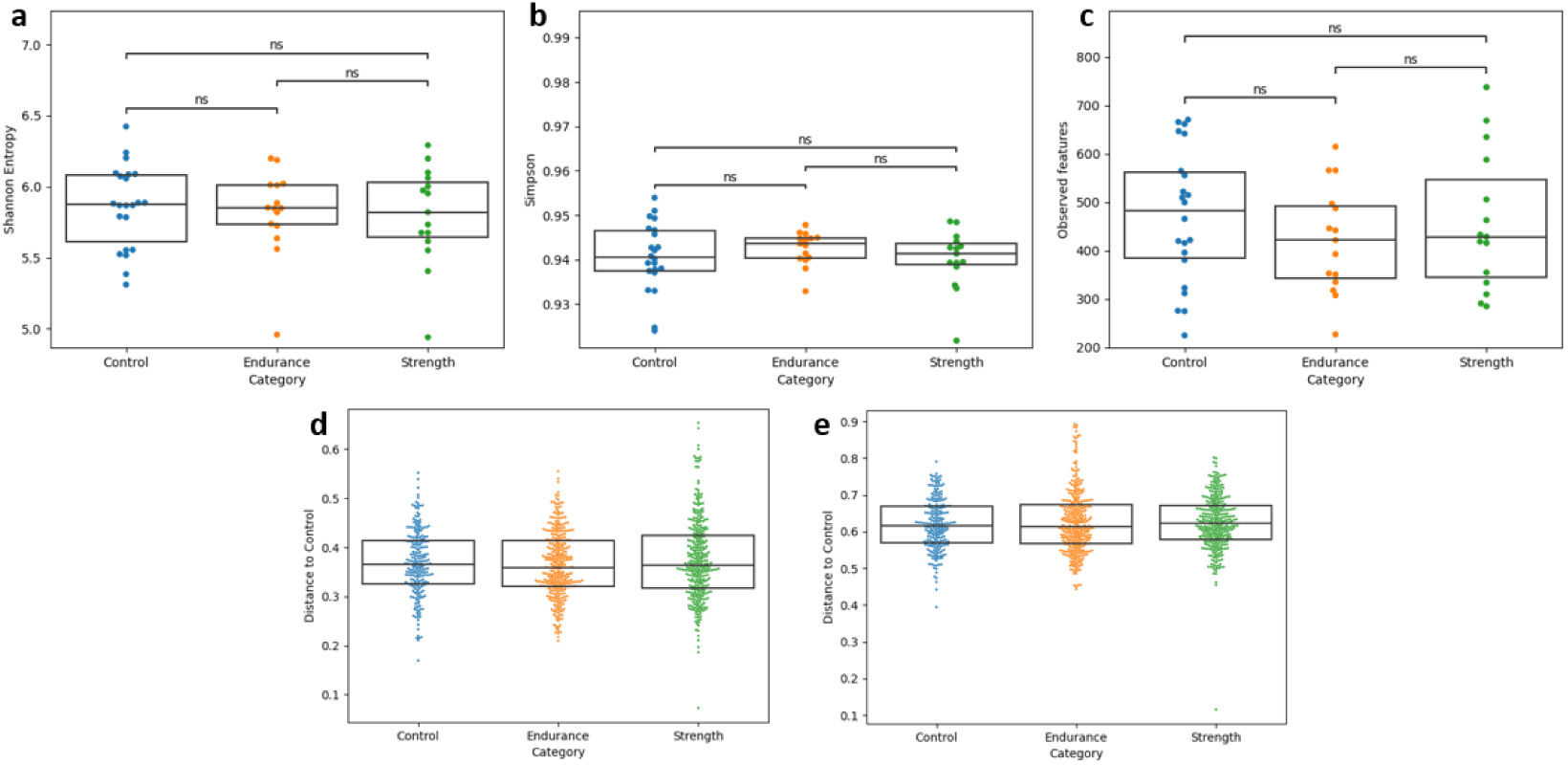
Comparison of alpha and beta diversity of the Control, Endurance and Strength groups. **a)** Shannon entropy. **b)** Simpson index. **c)** Observed features. **d)** Bray-Curtis distance. **e)** Jaccard distance.

There were no observable differences in the proportions of phyla among the control, strength and endurance groups (Figure 2). The most abundant genus was *Firmicutes*, followed by *Bacteroidetes* and *Actinobacteria* (average relative abundance of 66.7%, 25.9%, and 5.9%, respectively). *Bacteroidetes* contributed to approximately half of the population in the three samples (one control and two strength), whereas *Firmicutes* constituted the majority in most individuals. We investigated enterotypes, which were previously defined as microbiota profile types [28], also applied to athletic populations (with *Bacteroides*-driven enterotype being associated with high fat and protein intake of strength athletes while *Prevotella*-driven enterotype linked to simple carbohydrates commonly ingested by endurance athletes [29]. *Bacteroides*-driven and *Ruminococcus*-driven enterotypes were equally prevalent in the control group (40.9% each). *Bacteroides*-driven enterotype was dominant in the endurance group (46.7%), whereas *Prevotella*-driven enterotype was dominant in the strength group (50.0%).

**Figure 2.A.**
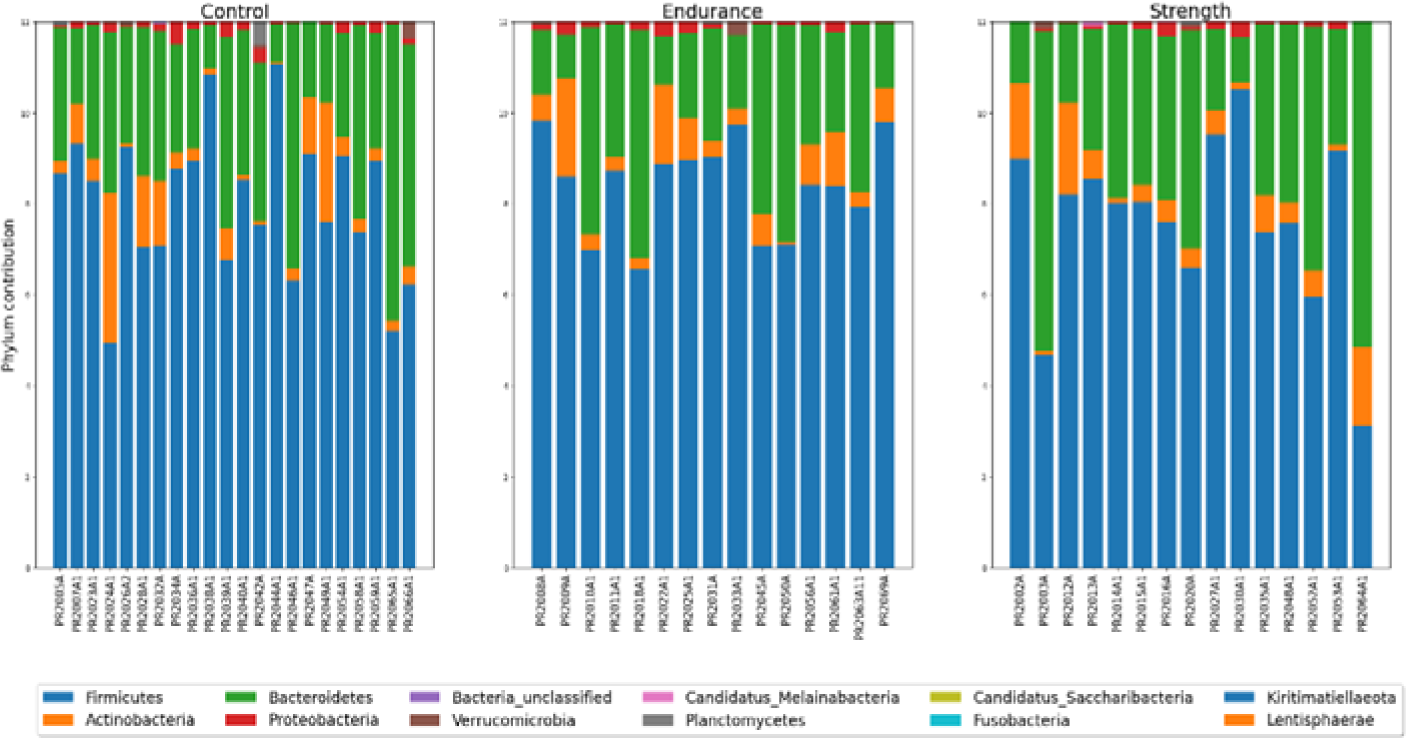
Phyla contributions in every sample, grouped by Control, Endurance and Strength. **B.** Enterotype contributions to Control, Endurance and Strength groups

**Figure 2.B.**
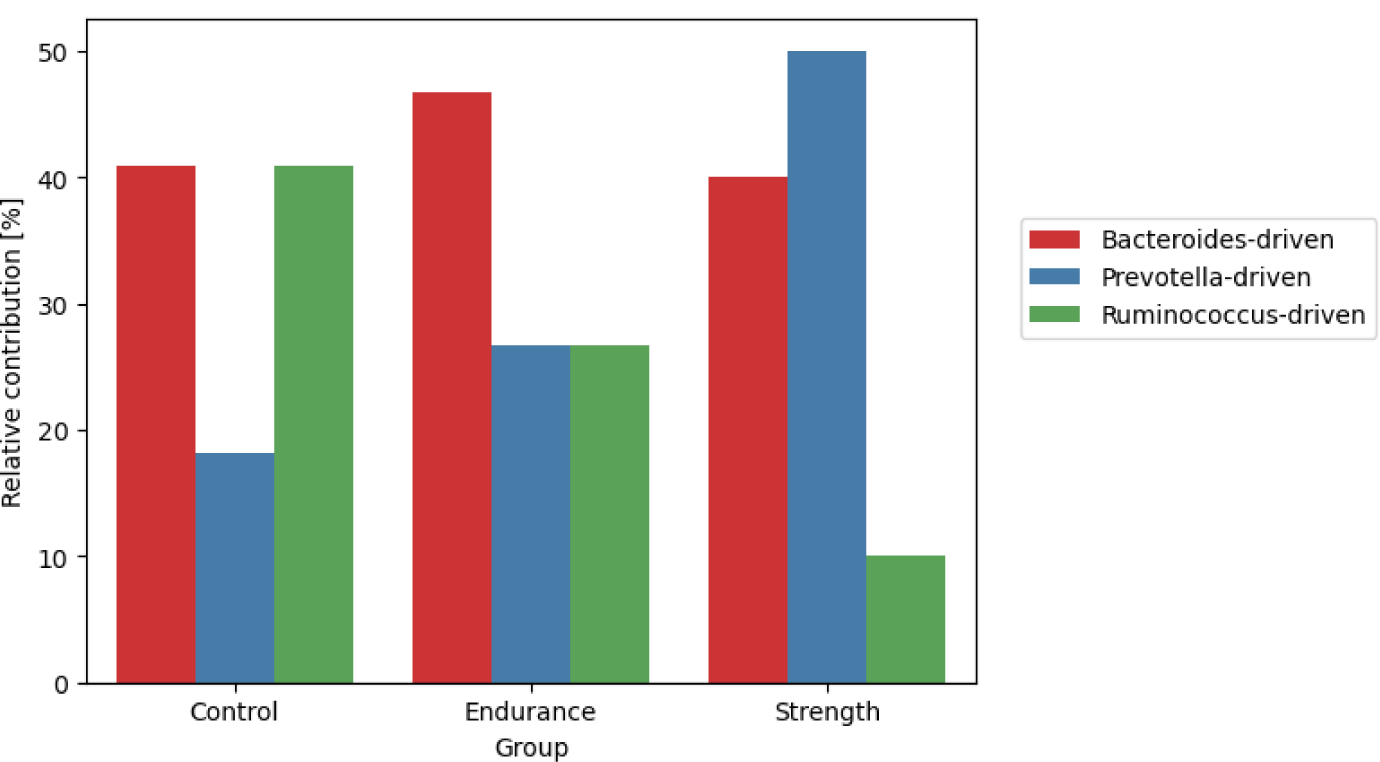
Enterotype contributions to Control, Endurance and Strength groups

The intersection of species and functions identified in each group is presented in Figure 3. While nearly 34% of all species identified were unique to either group, only 7% of the functions occurred in a single group. An equal number of species were shared between control and endurance and control and strength, whereas three times fewer species were shared between endurance and strength. We were particularly excited to see an enrichment of indigestible carbohydrate degrading bacteria [30] (*Blautia genus*) in the endurance group, suggesting performance-specific adaptations. We also observed a presence of *Christensenella minuta* (a potential therapeutic target associated with the health of the gut) [31] and *Gordonibacter urolithinfaciens* (one of the few species able to convert ellagic acid to urolithins) [32]. Furthermore, a number of probiotic taxa (*Lactobacillus kalixensis* [33], and *Leuconostoc pseudomesenteroides* [34], *Monoglobus pectinilyticus* [35]) with yet poorly defined mechanisms of action were found in the trained and not in the non-trained individuals. Control shared more pathways with endurance, whereas a similar number of functions were shared between control and strength and strength and endurance. The complete list of species and functions unique to each group is shown in Supplementary Table 3.

**Figure 3.**
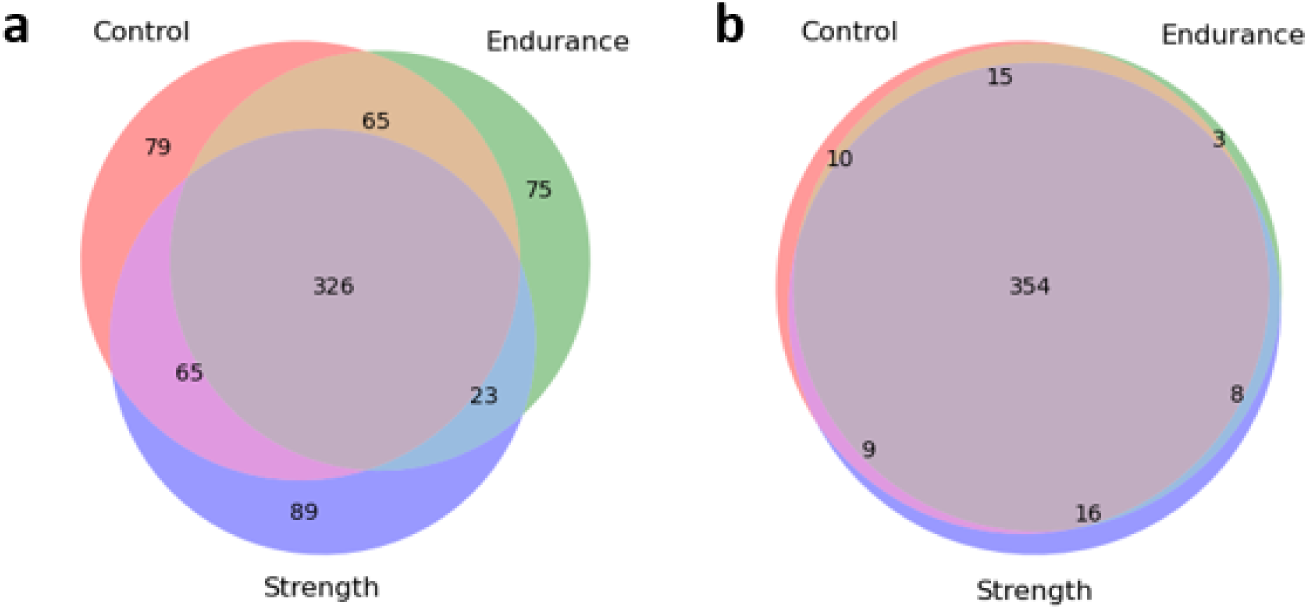
Overlap of a) species and b) functions identified in each group at baseline.

### 3.2. Comparison of microbiome alterations due to sports interventions in different groups

Although no major differences between groups were identified at baseline, distinct microbiome responses to the two types of interventions (Wingate Anaerobic Test and Bruce Trademill Test) were expected.

The initial step of the analysis involved comparing the strength, endurance and control groups cross-sectionally at each time point (W1, W2, B0, B1, and B2). Differential abundance analysis with ANCOM showed no significant taxonomic, functional, or gene differences between the control, strength and endurance groups. The results produced by LEfSe, which is known to have less stringent thresholds than ANCOM [36], also did not reveal many variations (Table 2). Identified as significantly more abundant in endurance versus control and strength at most time points was *Blautia sp AF19 10LB*, though not yet known to have implications in human physiology, but has shown a series of potential probiotic properties [37]. No other species were identified at multiple time points within a single group.

**Table 2.** Species identified as specific per group after being identified as significantly more abundant in the group of interest in comparison to the other two groups by LEfSe.

| Timepoint | Control-specific | Endurance-specific | Strength-specific |
| --- | --- | --- | --- |
| B0 | - | *Blautia sp AF19 10LB*<br>*Dorea longicatena* | *Catenibacterium sp AM22 15*<br>*Clostridium phooceensis* |
| W1 | *Bacteroides faecis* | *Fusicatenibacter saccharivorans* | *Ruminococcus sp AF41 9*<br>*GGB3571 SGB4778*<br>*Rothia mucilaginosa*<br>*Coprococcus catus* |
| W2 | *Phocaeicola massiliensis* | *Blautia sp AF19 10LB* | *GGB2977 SGB3959* |
| B1 | - | *Blautia sp AF19 10LB*<br>*Lacticaseibacillus paracasei* | *Clostridiales bacterium* |
| B2 | - | *Blautia sp AF19 10LB* | *Blautia sp MSK 20 85* |

B0: in the morning fasting before the Bruce Trademill Test, B1: the same day after the Bruce Trademill Test, B2: morning fasting after the Bruce Trademill Test on an empty stomach, W1: the same day after the Wingate Anaerobic Test, W2: morning fasting after the WAnT on an empty stomach

LEfSe analysis of functional profiles indicated that one pathway (PWY 5695: inosine 5’-phosphate degradation) was significantly more enriched in endurance versus control, but not in strength. Six pathways were identified in endurance versus strength, but not in the control. These included PWY 7229: superpathway of *de novo* adenosine nucleotide biosynthesis, PWY 841: superpathway of *de novo* purine nucleotide biosynthesis, PWY 7400: L-arginine biosynthesis, PWY 6126: superpathway of *de novo* adenosine nucleotide biosynthesis, PWY 6703: preQ0 biosynthesis, and PWY 7228: superpathway of *de novo* guanosine nucleotide biosynthesis. Interestingly, the same pathways have previously been observed as enriched in elite Irish athletes performing high static component sports such as triathlon or cycling [38]. The six pathways enriched in endurance versus strength but not control particularly attracted our attention, as they could indicate mechanisms somehow compromised as a result of strength training. Indeed, being linked to NO synthesis (resulting in improved mitochondrial biogenesis and gas exchange) and influencing the TCA cycle, they appeared to be more aerobic-oriented, though the effects of amino acids such as L-arginine had been positively linked to both aerobic and anaerobic performances [39]. No significantly enriched genes were identified in this study. No significant genes were identified in this study.

Longitudinal analysis of alpha (Shannon entropy) and beta (Bray-Curtis) diversities revealed no significant differences between any pair of the Bruce intervention timepoints (T0-T1, T0-T2, T1-T2), within or between any of the groups (data not shown). Due to the lack of baseline Wingate samples, longitudinal analysis of the anaerobic intervention could not be performed. However, as baseline Bruce and WAnT samples appeared to not be significantly distant from one another (further explanation in the Methods section), we decided to substitute the missing WAnT samples with the baseline Bruce samples to get a general overview of the changes linked to the Wingate intervention. Interestingly, *Alistipes communis*, yet underexplored regarding its implications for human health [40] but previously positively correlated with resistance training [41], occurred to change markedly throughout the WAnT intervention. Despite a slight net decrease across all individuals combined (average net change = − 0.0004), it appeared to increase steadily in the strength group (Figure 4). Statistical analysis confirmed a significant decrease in *Alistipes communis* in the control (Wilcoxon signed-rank test, FDR P-value = 0.03) and endurance (Wilcoxon signed-rank test, FDR P-value = 0.04) between the baseline and the second time point post-Wingate intervention. Furthermore, there was a statistically significant difference between strength and control (Mann-Whitney U test, FDR P-value < 0.02) and between strength and endurance (Mann-Whitney U test, FDR P-value < 0.02).

**Figure 4.**
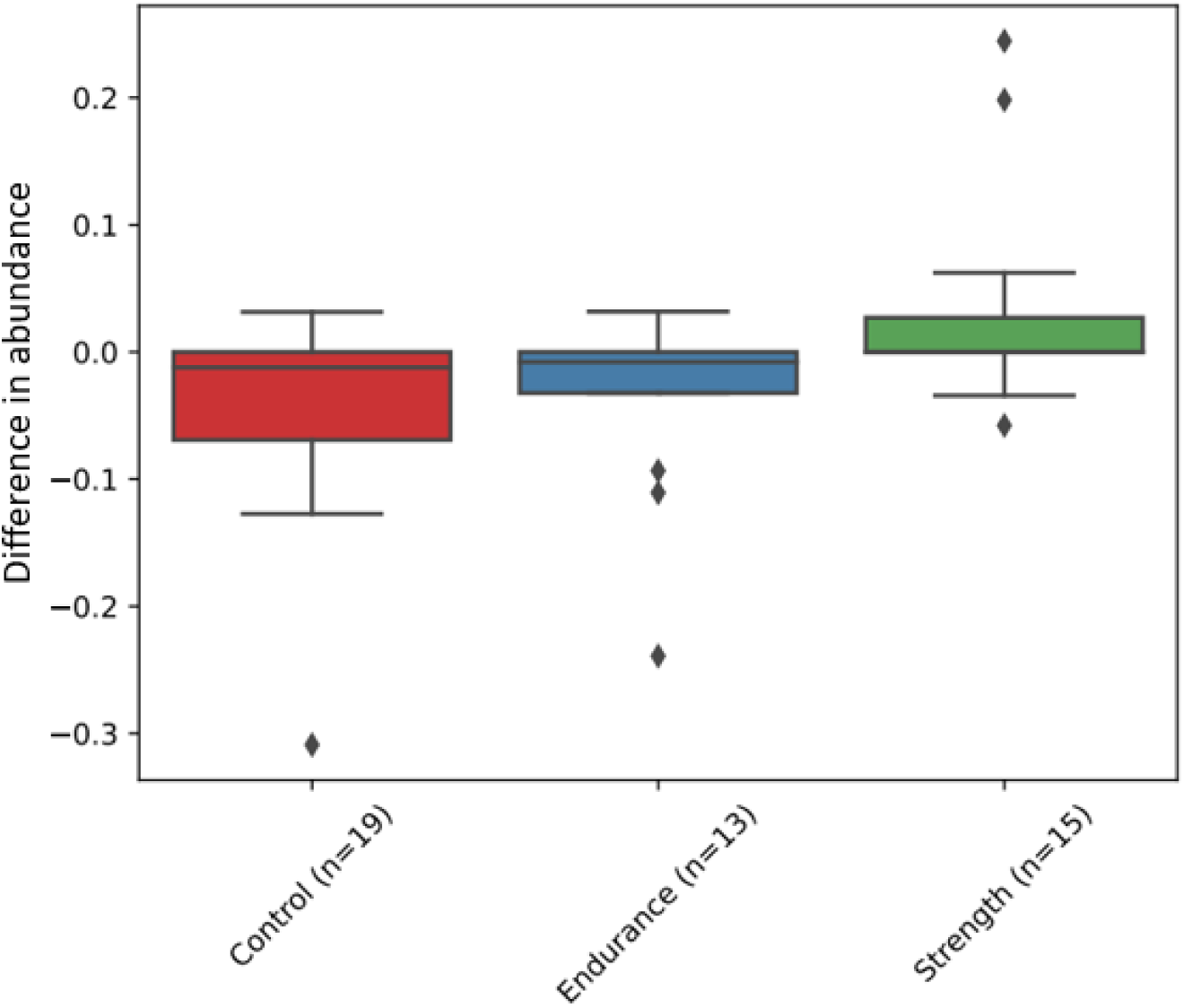
The difference in abundance of *Alistipes communis* between baseline (in the morning fasting before the WAnT) and W2 (the following day after the WAnT on an empty stomach) per category.

88 species were identified throughout the Bruce Test intervention. *Blautia wexlerae, Bacteroides ovatus* and *Bacteroides thetaiotaomicron* displayed the greatest positive net average changes (average net changes of 0.007, 0.002, and 0.002, respectively). *Blautia wexlerae* is a short-chain fatty acid producer [42], while *Bacteroidetes* have a wide range of genes responsible for carbohydrate metabolism [43]. The most significant net negative change was experienced by *Fusicatenibacter saccharivorans* (average net change of -0.009), which is also a short-chain fatty acid producer and a core component of the human gut microbiota [44].

Several species showed different responses across groups. *Anaeroburyticium hallii*, a strict anaerobe that utilizes glucose and is involved in acetate and lactate metabolism to butyrate and hydrogen [45], experienced an initial increase at B1 and a subsequent decrease at B2 in control, but a reverse trend in endurance and strength. *Ruminococcus bromii* showed an initial decrease followed by an increase at B2 in strength and endurance, but no change in the control. It is a known short-chain fatty acid producer with anti-inflammatory properties [46]. *Bacteroides xylanisolvens*, associated with its probiotic properties [47], displayed a gradual increase in all groups at first, but a strong decrease in the control at B2, while it continued to increase in strength and endurance. Finally, *Roseburia hominis* experienced an increase in strength and endurance but not in control. This short-chain fatty acid producer, involved in maintaining homeostasis in the gut [48], has previously been shown to increase in abundance in well-trained rowers throughout an ultra-endurance event [49]. Another species that increased in the rowing event was *Dorea longicatena*, which also showed expansion in endurance and control at B1 and strength at B2.

### 3.3. Species linked to different aspects of sports performance

Despite the interesting trends identified by Qiime2 in the longitudinal analysis, each group contained substantial variation. Therefore, it is not appropriate to define these findings as group-describing characteristics. Because the differences between the groups were not as significant as expected, we wondered about the actual physical variation between and within groups and whether it could explain the diversity in microbiome shifts due to the applied interventions. The features considered included general fitness (fitness score), aerobic capacity (VO_2_max measured during the Bruce test), anaerobic capacity (maximum and average power per kilogram of body weight during the Wingate test), and the ability to maintain power (fraction of max and average power per kilogram of body weight maintained in the second part of the WAnT). Analysis of the WAnT and Bruce intervention results uncovered a lack of visible differences between the groups regarding fitness parameters (Supplementary Figure 1).

Furthermore, significant variations within the groups were present, and some even resembled bimodal distributions (VO_2_max in endurance, Average power per body weight for strength). This could explain the wide range of longitudinal taxonomic trends across individuals from the same group described in the earlier sections. No visible correlations of enterotypes with fitness parameter values were observed, apart from two endurance and strength individuals with seemingly better abilities to maintain both average and maximum power throughout the Wingate intervention, who were enterotype *Ruminococcus* dominant enterotypes.

MaAsLin was used to search for linear correlations between the microbiota at baseline and parameter values (data not shown). The lack of significant output hits could be explained by either an absence of interactions or more complex connections of bacterial species with sports performance testing.

MaAsLin was used to search for linear correlations between the microbiota at baseline and parameter values (data not shown). The lack of significant output hits could be explained by either an absence of interactions or more complex connections of bacterial species with sports performance testing. When we applied the SparCC method, a range of microbial, functional, and gene associations with the fitness score, VO_2_max, average power, and maximal power were identified (Figure 5), suggesting the latter. The species with the highest correlation values were associated with VO_2_max. These included *Bifidobacterium adolescentis* (r=0.18) and *Bifidobacterium longum* (r=0.14), which are popular probiotics with positive effects on athletic performance [50,51]. We also found strong associations between *Phocaeicola vulgatus* and *Roseburia intestinalis* with fitness score (r values of 0.17 and -0.17, respectively). Although the function of *Phocaeicola vulgatus* remains ambiguous [52], *Roseburia intestinalis* is known to prevent inflammation and maintain homeostasis [53]. The correlations were specific to the parameters, and no universal set of bacteria influencing all parameters could be identified in the same way. A complete list of the top species correlated with each fitness parameter is shown in Supplementary Table 5.

**Figure 5.**
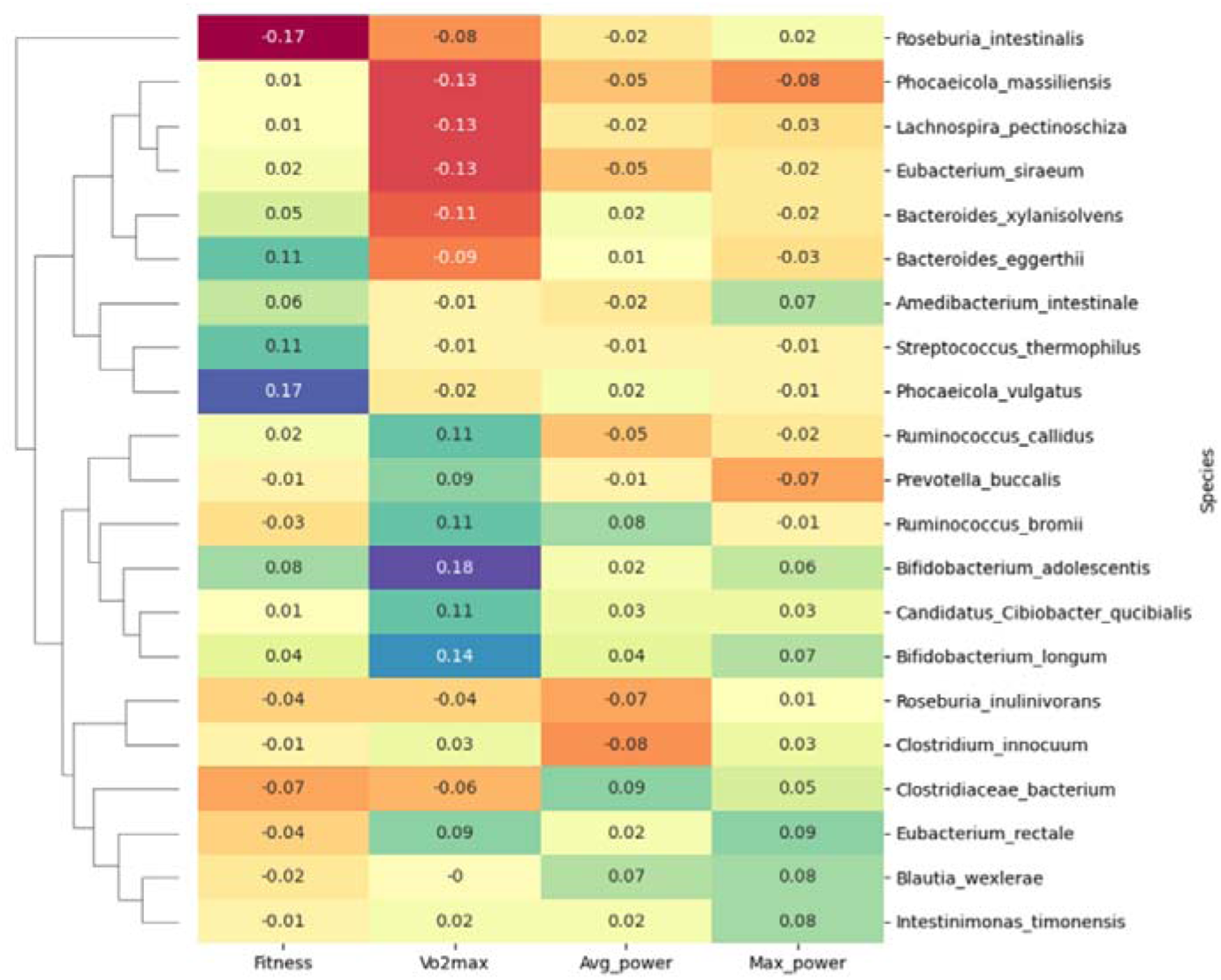
Top SparCC correlations of species with fitness parameters.

When we investigated top SparCC correlations using functions instead of taxa (r >= 0.6), only *PWY-7238: sucrose biosynthesis II* correlated with two different parameters (r ≥ 0.6, fitness score and average power). However, the direction of the correlations was the opposite (positive for fitness score and negative for average power), and connections with different species were noted (*Bifidobacterium adolescentis* for fitness score and *Blautia wexlerae* for average power). The functions with the highest correlations included inosine 5’-phosphate degradation and phosphopantothenate biosynthesis for fitness score, purine ribonucleoside degradation and chorismite biosynthesis for VO_2_max, sucrose biosynthesis and L-arginine biosynthesis for average power, and molybdopterin biosynthesis and fatty acid biosynthesis for maximum power. Interestingly, some species contributing to the functions were present in association with multiple parameters, and the species shown in Figure 6 were not among them. *Fusicatenibacter saccharivorans* was associated with functions correlated with fitness score, VO_2_max, and maximum power; *Blautia obeum* with VO_2_max and maximum power; *Faecalibacterium prausnitzii* with VO_2_max and average power; and *Ruminococcus torques* with fitness score and VO_2_max.

**Figure 6.**
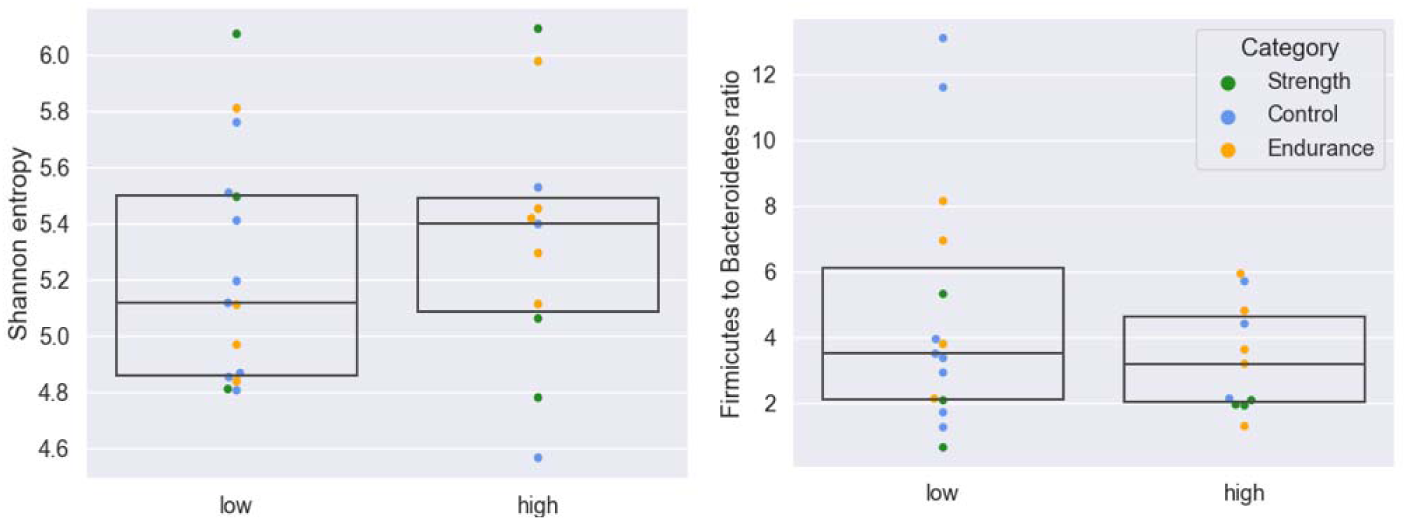
Comparison of high and low VO_2_max groups.

Analysis of gene correlations with fitness parameters revealed a few pathways correlated with multiple parameters, but usually in opposite directions, and associated with different species. RXN-9536 (3-oxo-meristoyl-[acyl-carrier protein] reductase), however, was associated with the fitness score and average power and positively correlated with *Blautia obeum* in both cases.

In addition to the microbiota correlations with parameter values, the differences between the relatively high and low parameter groups were investigated. VO_2_max and average power per kilogram were selected as the best representatives of anaerobic (WAnT) and aerobic (Bruce Test) capacities. Naturally, because all participants were physically active, fit and healthy, both average power and VO_2_max values were substantially higher than in the general population (“Very good“ and “Excellent,” according to VO_2_max reference values) [54]. Therefore, we decided not to adhere to the standard VO_2_max classification thresholds and based our comparison on the distribution of values in our data, in effect comparing good but relatively high and relatively low VO_2_max and average power values across participants. Ultimately, two groups per parameter were established (relatively low average power: < 8 W/kg and VO_2_max < 55 ml/kg/min, and relatively high power: > 8.5 W/kg, and VO_2_max > 62 ml/kg/min), further referred to as high and low groups. Participants with intermediate values were not considered in the subsequent comparisons.

For the high and low average power per kilogram groups, no alpha or beta diversity differences were observed. According to ANCOM, *Bifidobacterium longum* and *Bifidobacterium adolescentis* were the most differentially abundant species in the high group (CLR values above 0.4), whereas *Prevotella copri clade A* and *Ruminococcaceae unclassified* in the low group (CLR values above 0.3). However, these differences were not statistically significant.

Comparison of high and low VO_2_max groups revealed subtle differences in alpha diversity (Figure 7, left), but no variation in beta diversity (data not shown). *Firmicutes* to *Bacteroides* ratio, associated with VO_2_max [55], appeared to be more significant in some samples in the low versus high group (Figure 6, right). Differential abundance with ANCOM did not yield statistically significant results but revealed species with the greatest differential enrichment. These included *Bidifobacterium longum*, *Candidatus Cibiobacter qucibialis* and *Bifidobacterium adolescentis* in the high VO_2_max group (CLR values above 0.5), and *Roseburia intestinalis*, *Prevotella copri clade A, Lachnospira pectinoschiza* in the low VO_2_max group (CLR values nearly 0.4 and above).

**Figure 7.**
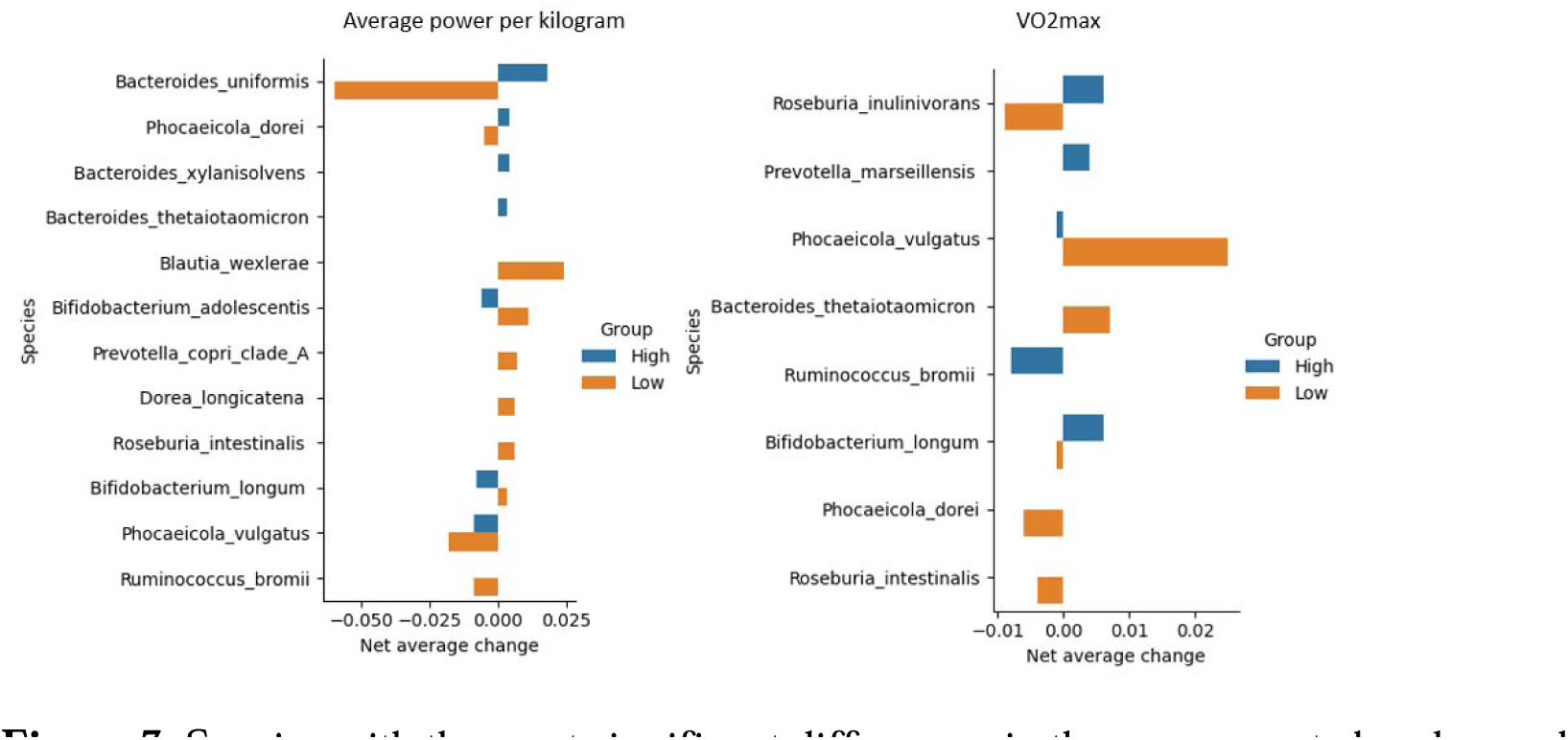
Species with the most significant differences in the average net abundance change between high and low average power and VO_2_max groups.

Longitudinal analyses identified the species with the most remarkable net average changes throughout the Bruce intervention. Figure 7 shows the top taxa with different high- and low-parameter group patterns. The most significant discrepancy in the average power group was *Bacteroides uniformis*, which drastically decreased in abundance in the low power group and increased in the high power group. *Bacteroides uniformis* facilitates glucose production and enhances physical performance [56]. The VO_2_max was *Roseburia inulinivorans*, with a slightly more significant decrease in the low group than in the high group. Like other species from the *Roseburia* genus, *Roseburia inulinivorans* is a short-chain fatty acid producer [57].

Interestingly, opposite trends were noted when comparing species associated with the high or low groups in the average power and VO_2_max categories. *Bifidobacterium longum* increased in the high group and decreased in the low group in terms of VO_2_max and experienced a reverse shift in the average power category. *Phocaeicola vulgatus* decreased in abundance in both high groups and increased in VO_2_max, but decreased in average power. Finally, while no longitudinal changes were observed in the high groups regarding *Roseburia intestinalis* abundance, it was enriched in the low average power group and became less abundant in the low VO_2_max group.

Finally, the bimodal distribution identified within groups in the maximum strength per kilogram plot (Figure 5) allowed us to fully differentiate between individuals with different training statuses and physiological characteristics. As a result, although small, two contrasting groups were established: strength individuals with visibly higher maximum average strength than the rest of the group and less trained individuals from the control group (Figure 8).

**Figure 8.**
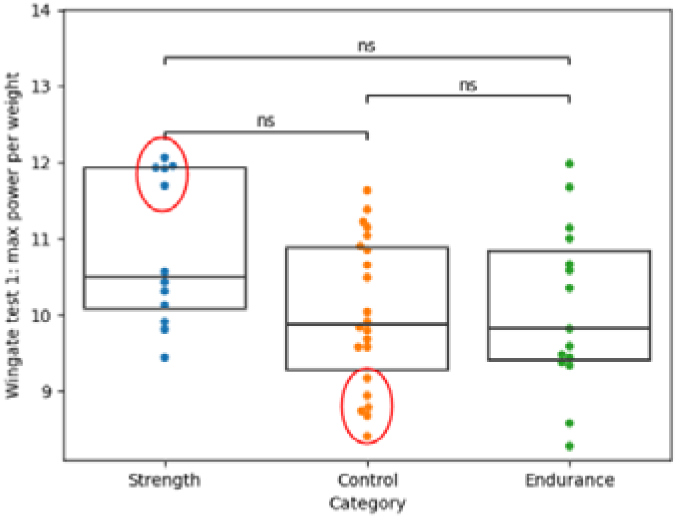
Distributions of maximum power values in Strength, Control and Endurance groups, with marked high-power Strength and low-power Control individuals.

No alpha or beta diversity differences between the groups were identified (data not shown). In terms of differential abundance with ANCOM, although no significant results were identified, there were substantially more abundant species in either group. The species most enriched in the low-power control group were *Ruminococcus bicirculans, Bacteroides ovatus, Phocaeicola massiliensis* and *Bacteroides caccae* (CLR values above 0.5). In the high power strength group, *Lachnospira eligens, Faecalibacterium prausnitzii, Bacteroides stercoris, Bifidobacterium pseudocatenulatum* and *Phocaeicola plebeius* were more enriched (CLR values above 0.5). Longitudinal analysis of the Wingate intervention revealed a significant increase in the abundance of *Dysosmobacter welbionis* in the high group (average net change of 0.001) and a decrease in this species in the low group (average net change, 0.001). *Dysosmobacter welbionis* is found in approximately 65% of healthy individuals and is negatively correlated with TBMI and fasting blood glucose levels in obese individuals with metabolic syndrome [58]. No other volatile features were observed in the high-power group. The most significant increase in the low group was observed for *Bifidobacterium adolescentis* and a decrease in *Eubacterium rectale* (average net changes of 0.018 and –0.041, respectively).

## 4. Discussion

Most recent studies have highlighted that physical exercise positively influences the diversity and composition of the gut microbiome [59,60]. Moreover, increased maximal oxygen uptake is also positively correlated with microbial diversity, which can be especially noticeable when comparing sedentary individuals with elite athletes, for whom this value can be twice as high [61]. To the best of our knowledge, no study has yet been conducted on the microbiome of physically active people but not training compared to those who train.

Therefore, in our study, we compared microbiome alterations of non-trained (control) to trained (strength, endurance) physically active individuals as a result of two types of maximal intensity interventions (Wingate Anaerobic Test and Bruce Treadmill Test) and identified sport-level and training-level microbiome characteristics. We discovered similar microbiome features across all participants, which was not surprising considering that all individuals were healthy, fit, and physically active. However, differences that we discovered, especially those linked to specific aspects of performance, have previously been found in elite athletes. We were particularly excited to see an enrichment of indigestible carbohydrate degrading bacteria in the endurance group, suggesting performance-specific adaptations. Furthermore, a number of probiotic taxa with yet poorly defined mechanisms of action were found in the trained and not in the non-trained individuals.

In our study, we also observed a positive influence of physical fitness on microbial diversity and its correspondence with VO_2_max, but not between control, strength and endurance, probably because they were all physically active. This confirms that physical activity may be a hallmark of microbiome diversity, which was observed to correlate with general health and homeostasis maintenance [62].

Several species were consistently identified throughout the analysis. We were particularly interested in *Bifidobacterium longum* and *Bifidobacterium adolescentis*, both positively correlated with all fitness parameters (robust correlations for VO_2_max) and an increase in abundance in the high VO_2_max group throughout the Bruce Test intervention. These two probiotic species are well represented in commercial products [63]. However, the effects of probiotic products on athletic performance are still poorly understood [64]. While we did not determine strain-level taxonomy from our samples [65], it was interesting that our results supported the beneficial effects of *Bifidobacterium longum* and *Bifidobacterium adolescentis* on general fitness and aerobic capacity.

In contrast, we noted negative correlations between *Bacteroides* species and VO_2_max. Although *Bacteroides* are generally assumed to be present in a healthy microbiome [66], *Firmicutes* to *Bacteroidetes* ratio has previously been associated with a high VO_2_max [55]. While we did not observe significant correlations between *Firmicutes* to *Bacteroides* ratio and VO_2_max, the negative correlation of numerous *Bacteroides* species with VO_2_max was interesting. Furthermore, we discovered a positive correlation between *Prevotella* and VO_2_max, and a high *Prevotella* to *Bacteroides* ratio is known to be associated with improved glucose metabolism [67] and increased glycogen storage. A high *Prevotella* to *Bacteroides* ratio was also previously observed in top Polish endurance athletes [68], but did not differ among e-sport players and physical education students ^64^.

Furthermore, we observed positive correlations between common SCFA producers (*Blautia wexlerae, Eubacterium rectale* and *Intestinimonas timonensis*) and maximal power during the WAnT intervention. SCFAs can be used as additional substrate for metabolism [69], usually desired in endurance sports. However, butyrate also induces beneficial alterations in skeletal tissues [70], which could explain the correlation between butyrate producers and power.

The main limitation of our study was its small sample size and missing W0 data This limitation could affect the interpretation of the results. However, since the baseline samples of the Bruce and WAnT tests did not show significant differences, we replaced the data form missing WAnT samples with the baseline Bruce samples. This substitution allowed us to obtain a comprehensive overview of the changes associated with the WAnT intervention. While visible differences between groups of individuals could be identified, the trends were highly individualized. A study with a larger sample size could potentially identify within-group trends and perhaps enable the determination of aspects related to responses to a particular type of intervention. Furthermore, we should focus mainly on individuals who compete at an international level in their sports disciplines. This would provide an opportunity to examine sport-specific microbiome associations only for individuals who are well adapted to aerobic or anaerobic efforts, which is the primary association with their discipline.

Moreover, it is essential to remember that such highly sport-specific physical efforts always affect nutritional factors and dietary requirements (the proportion of proteins, fatty acids, carbohydrates, vitamins, and specific supplements), which may influence microbiome homeostasis. Additionally, we should try to examine people who are not physically active and do not attend even occasional physical activities. This provides an opportunity to determine how the microbiome is regulated without physical activity.

Overall, we demonstrated that healthy and fit participants, regardless of their training level, displayed similar features to a healthy microbiome. However, we observed various individual responses, suggesting complex microbiota interactions or confounding factors. In light of the absence of W0 samples, we recommend conducting further research that includes the inclusion of those samples. Therefore, we suggest further research with more participants in shared and individualized responses, both at the species and functional levels. Including more extreme cases, such as obese sedentary or elite athletes, would allow for better distinction between health-related and athlete-related microbiome features. Furthermore, it would also be beneficial to include additional data, such as metabolomics, that could provide more profound insights into the differences between individuals and their responses to interventions (at any tissue level). We believe that the proposed approaches could move the field closer to explaining the ambiguities of the current findings and revealing the global role of the microbiome in physically active populations.

## Acknowledgments

We sincerely thank the members of the Sanprobi sp. z o.o. sp. k, Szczecin in Poland, especially Karolina Skonieczna-Żydecka, Anna Wierzbicka-Woś, Igor Łoniewski and Mariusz Kaczmarczyk for their cooperation during the validation procedure of microbiome library preparation.

## Data availability statement

The data supporting the findings of this study are openly available in the European Nucleotide Archive (ENA) at https://www.ebi.ac.uk/ena/, reference number PRJEB60692.

## Institutional Review Board Statement

Approved by Bioethics Committee for Clinical Research of the Regional Medical Society in Gdansk (KB-27/18).

