## Supplementary materials for "Microbiome Features Associated with Performance Measures in Non-Athletic and Athletic Individuals"

### 1 Supplementary Figures and Tables

**Supplementary Figure 1.** Distributions of fitness parameter values per group with marked enterotypes. Green: *Bacteroides*-dominant, Blue: *Prevotella*-dominant, Orange: *Ruminococcus*-dominant.

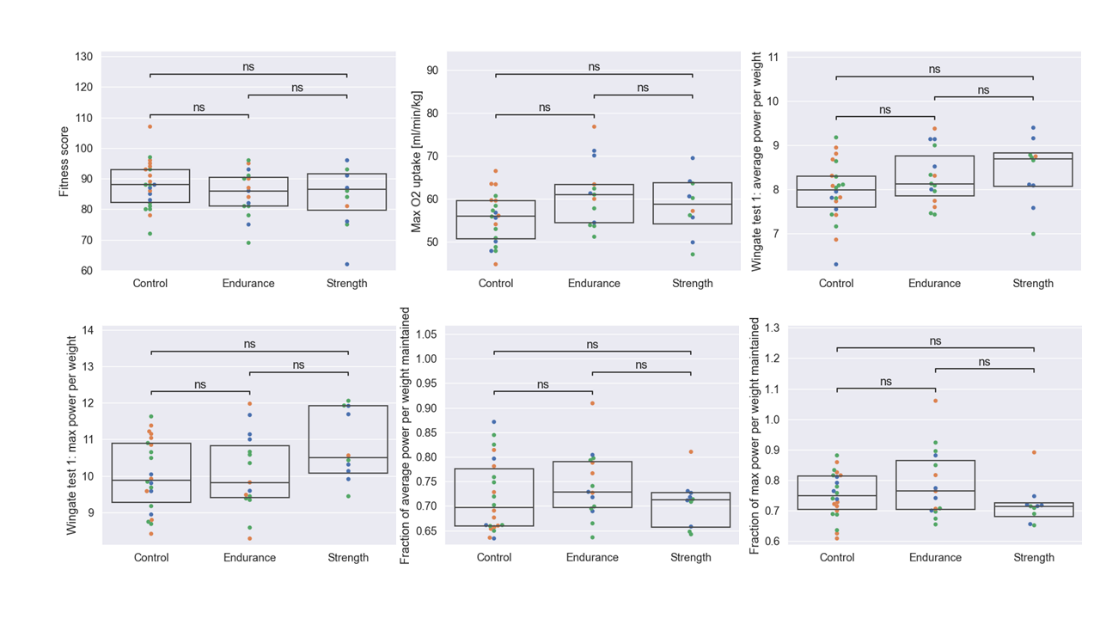

**Supplementary Table 1.** Alpha diversity results for Control, Endurance and Strength groups at baseline.

| **Shannon entropy** | | | | |
| --- | --- | --- | --- | --- |
| **Timepoint** | **Groups** | **Kruskal-Wallis H** | **p-value** | **q-value** |
| W1 | Control vs Strength | 0.33 | 0.57 | 0.57 |
|  | Control vs Endurance | 2.22 | 0.14 | 0.41 |
|  | Strength vs Endurance | 0.48 | 0.49 | 0.57 |
| W2 | Control vs Strength | 0.01 | 0.92 | 0.92 |
|  | Control vs Endurance | 0.19 | 0.67 | 0.92 |
|  | Strength vs Endurance | 0.07 | 0.79 | 0.92 |
| B0 | Control vs Strength | 0.28 | 0.60 | 0.76 |
|  | Control vs Endurance | 0.12 | 0.73 | 0.76 |
|  | Strength vs Endurance | 0.10 | 0.76 | 0.76 |
| B1 | Control vs Strength | 0.04 | 0.85 | 0.85 |
|  | Control vs Endurance | 0.56 | 0.46 | 0.68 |
|  | Strength vs Endurance | 0.57 | 0.45 | 0.68 |
| B2 | Control vs Strength | 0.78 | 0.38 | 0.61 |
|  | Control vs Endurance | 0.70 | 0.40 | 0.61 |
|  | Strength vs Endurance | 0.00 | 0.95 | 0.95 |
| **Simpson** | | | | |
| **Timepoint** | **Groups** | **Kruskal-Wallis H** | **p-value** | **q-value** |
| W1 | Control vs Strength | 0.43 | 0.51 | 0.77 |
|  | Control vs Endurance | 0.02 | 0.90 | 0.90 |
|  | Strength vs Endurance | 0.63 | 0.43 | 0.77 |
| W2 | Control vs Strength | 0.56 | 0.45 | 0.45 |
|  | Control vs Endurance | 2.22 | 0.14 | 0.31 |
|  | Strength vs Endurance | 1.51 | 0.22 | 0.33 |
| B0 | Control vs Strength | 0.80 | 0.37 | 0.55 |
|  | Control vs Endurance | 0.00 | 0.98 | 0.98 |
|  | Strength vs Endurance | 1.50 | 0.22 | 0.55 |
| B1 | Control vs Strength | 0.00 | 0.95 | 0.95 |
|  | Control vs Endurance | 0.66 | 0.42 | 0.65 |
|  | Strength vs Endurance | 0.61 | 0.44 | 0.65 |
| B2 | Control vs Strength | 0.26 | 0.61 | 0.61 |
|  | Control vs Endurance | 1.81 | 0.18 | 0.28 |
|  | Strength vs Endurance | 1.76 | 0.18 | 0.28 |
| **Observed features** | | | | |
| **Timepoint** | **Groups** | **Kruskal-Wallis H** | **p-value** | **q-value** |
| W1 | Control vs Strength | 0.43 | 0.51 | 0.51 |
|  | Control vs Endurance | 1.35 | 0.25 | 0.37 |
|  | Strength vs Endurance | 4.10 | 0.04 | 0.13 |
| W2 | Control vs Strength | 0.56 | 0.45 | 0.45 |
|  | Control vs Endurance | 2.22 | 0.14 | 0.31 |
|  | Strength vs Endurance | 1.51 | 0.22 | 0.33 |
| B0 | Control vs Strength | 1.07 | 0.30 | 0.72 |
|  | Control vs Endurance | 0.13 | 0.72 | 0.72 |
|  | Strength vs Endurance | 0.23 | 0.63 | 0.72 |
| B1 | Control vs Strength | 1.11 | 0.29 | 0.54 |
|  | Control vs Endurance | 0.37 | 0.54 | 0.54 |
|  | Strength vs Endurance | 0.78 | 0.38 | 0.54 |
| B2 | Control vs Strength | 0.17 | 0.68 | 0.68 |
|  | Control vs Endurance | 0.17 | 0.68 | 0.68 |
|  | Strength vs Endurance | 0.85 | 0.36 | 0.68 |

B0: in the morning fasting before the Bruce Trademill Test, B1: on the same day after the Bruce Trademill Test, B2: morning fasting after the Bruce Trademill Test on an empty stomach, W1: the same day after the Wingate Anaerobic Test, W2: morning fasting after the WAnT on an empty stomach

**Supplementary Table 2.** Beta diversity results for Control, Endurance and Strength groups at baseline.

| **Bray-Curtis** | | | | |
| --- | --- | --- | --- | --- |
| **Timepoint** | **Groups** | **Permanova pseudo-F** | **p-value** | **q-value** |
| W1 | Control vs Strength | 0.74 | 0.73 | 0.73 |
|  | Control vs Endurance | 0.79 | 0.69 | 0.73 |
|  | Strength vs Endurance | 1.40 | 0.14 | 0.42 |
| W2 | Control vs Strength | 0.62 | 0.89 | 0.89 |
|  | Control vs Endurance | 1.58 | 0.07 | 0.11 |
|  | Strength vs Endurance | 1.90 | 0.03 | 0.11 |
| B0 | Control vs Strength | 0.86 | 0.59 | 0.77 |
|  | Control vs Endurance | 0.71 | 0.77 | 0.77 |
|  | Strength vs Endurance | 1.28 | 0.22 | 0.65 |
| B1 | Control vs Strength | 0.83 | 0.65 | 0.65 |
|  | Control vs Endurance | 1.08 | 0.33 | 0.50 |
|  | Strength vs Endurance | 1.16 | 0.27 | 0.50 |
| B2 | Control vs Strength | 0.62 | 0.87 | 0.87 |
|  | Control vs Endurance | 0.65 | 0.85 | 0.87 |
|  | Strength vs Endurance | 0.78 | 0.65 | 0.87 |
| **Jaccard** | | | | |
| **Timepoint** | **Groups** | **Permanova pseudo-F** | **p-value** | **q-value** |
| W1 | Control vs Strength | 0.73 | 0.93 | 0.93 |
|  | Control vs Endurance | 1.00 | 0.41 | 0.62 |
|  | Strength vs Endurance | 1.27 | 0.09 | 0.28 |
| W2 | Control vs Strength | 0.84 | 0.74 | 0.76 |
|  | Control vs Endurance | 0.93 | 0.60 | 0.76 |
|  | Strength vs Endurance | 0.84 | 0.76 | 0.76 |
| B0 | Control vs Strength | 0.80 | 0.84 | 0.84 |
|  | Control vs Endurance | 0.95 | 0.51 | 0.84 |
|  | Strength vs Endurance | 0.82 | 0.81 | 0.84 |
| B1 | Control vs Strength | 1.02 | 0.37 | 0.55 |
|  | Control vs Endurance | 1.03 | 0.33 | 0.55 |
|  | Strength vs Endurance | 0.86 | 0.72 | 0.72 |
| B2 | Control vs Strength | 0.92 | 0.55 | 0.61 |
|  | Control vs Endurance | 0.90 | 0.61 | 0.61 |
|  | Strength vs Endurance | 0.96 | 0.46 | 0.61 |

B0: in the morning fasting before the Bruce Trademill Test, B1: on the same day after the Bruce Trademill Test, B2: morning fasting after the Bruce Trademill Test on an empty stomach, W1: the same day after the Wingate Anaerobic Test, W2: morning fasting after the WAnT on an empty stomach

**Supplementary Table 3.** Species and functions unique to Control, Endurance and Strength.

| **Group** | **Unique species** | **Unique functions** |
| --- | --- | --- |
| Control | *Actinomyces graevenitzii*  *Alistipes sp An31A*  *Bacilli unclassified SGB6428*  *Bacteroidaceae unclassified SGB1909*  *Bifidobacterium angulatum*  *Bifidobacterium pullorum*  *Butyricimonas SGB1783*  *Butyricimonas faecalis*  *Candidatus Allochristensenella caecavium*  *Candidatus Avispirillum faecium*  *Candidatus Parachristensenella avicola*  *Catabacter hongkongensis*  *Catenibacillus scindens*  *Clostridia unclassified SGB3983*  *Clostridia unclassified SGB6385*  *Clostridium perfringens*  *Clostridium saccharogumia*  *Collinsella massiliensis*  *Collinsella tanakaei*  *Coprobacillus cateniformis*  *Drancourtella sp An57*  *Eggerthella sp YY7918*  *Firmicutes bacterium AM41 11*  *GGB1256 SGB1684*  *GGB13431 SGB20700*  *GGB1421 SGB1958*  *GGB1632 SGB2240*  *GGB1689 SGB2321*  *GGB18381 SGB27144*  *GGB3034 SGB4030*  *GGB3118 SGB4130*  *GGB3474 SGB4637*  *GGB3532 SGB4720*  *GGB3677 SGB4990*  *GGB4239 SGB5731*  *GGB4254 SGB5782*  *GGB4482 SGB6176*  *GGB4482 SGB6177*  *GGB4616 SGB6391*  *GGB4713 SGB6526*  *GGB4750 SGB6579*  *GGB4770 SGB6604*  *GGB4964 SGB6927*  *GGB6544 SGB9242*  *GGB6546 SGB9245*  *GGB8965 SGB13825*  *GGB9186 SGB14125*  *GGB9288 SGB14243*  *GGB9358 SGB14333*  *GGB9434 SGB14812*  *GGB9523 SGB14922*  *GGB9551 SGB14958*  *GGB9559 SGB14968*  *GGB9623 SGB15076*  *GGB9668 SGB15165*  *GGB9694 SGB15201*  *GGB9709 SGB15238*  *Holdemania sp Marseille P2844*  *Intestinimonas massiliensis*  *Kiritimatiellae bacterium*  *Lachnoclostridium edouardi*  *Latilactobacillus curvatus*  *Lentisphaeria bacterium*  *Leuconostoc mesenteroides*  *Ligilactobacillus salivarius*  *Massilistercora timonensis*  *Mediterranea massiliensis*  *Megasphaera elsdenii*  *Ningiella SGB48271*  *Olsenella provencensis*  *Oscillibacter SGB15077*  *Prevotella SGB1589*  *Prevotella hominis*  *Prevotella lascolaii*  *Raoultella ornithinolytica*  *Senegalimassilia faecalis*  *Solobacterium SGB6833*  *Streptococcus lutetiensis*  *Trueperella pyogenes* | - ARGDEG-PWY:   superpathway of L-arginine. putrescine  and 4-aminobutanoate degradation   - ORNARGDEG-PWY:   superpathway of L-arginine and  L-ornithine degradation   - P163-PWY:   L-lysine fermentation to acetate and butanoate   - PWY-5464:   superpathway of cytosolic glycolysis (plants)  pyruvate dehydrogenase and TCA cycle   - PWY-6167:   flavin biosynthesis II (archaea)   - PWY-6328:   L-lysine degradation X   - PWY-6396:   superpathway of 2.3-butanediol biosynthesis   - PWY0-1221:   putrescine degradation II   - PWY0-41:   allantoin degradation IV (anaerobic)   - URSIN-PWY:   ureide biosynthesis |
| Strength | *Acidaminococcus intestini*  *Actinomyces SGB17168*  *Aggregatibacter segnis*  *Akkermansia sp KLE1605*  *Bacilli unclassified SGB6571*  *Bacteroidales unclassified SGB2076*  *Butyrivibrio crossotus*  *Candidatus Gastranaerophilales unclassified*  *SGB8630*  *Christensenella massiliensis*  *Clostridia unclassified SGB13972*  *Clostridia unclassified SGB14196*  *Clostridia unclassified SGB4372*  *Clostridia unclassified SGB6293*  *Clostridia unclassified SGB6344*  *Clostridiaceae bacterium NSJ 31*  *Clostridiales bacterium Choco116*  *Clostridium mediterraneense*  *Collinsella SGB14747*  *Eisenbergiella massiliensis*  *Emergencia timonensis*  *Eubacterium sp AF22 8LB*  *Faecalibacterium SGB15345*  *Flavonifractor sp An135*  *GGB1154 SGB1482*  *GGB1235 SGB1614*  *GGB1239 SGB1657*  *GGB1247 SGB1668*  *GGB1250 SGB1672*  *GGB1250 SGB1673*  *GGB13253 SGB20482*  *GGB1380 SGB1883*  *GGB1457 SGB2020*  *GGB1543 SGB2126*  *GGB1567 SGB2154*  *GGB1582 SGB2173*  *GGB1629 SGB2237*  *GGB3139 SGB4152*  *GGB32463 SGB47515*  *GGB3264 SGB4312*  *GGB3343 SGB4423*  *GGB3491 SGB4665*  *GGB3523 SGB4702*  *GGB3637 SGB4930*  *GGB4266 SGB5809*  *GGB4542 SGB6263*  *GGB45620 SGB63333*  *GGB4590 SGB6349*  *GGB4609 SGB6383*  *GGB4672 SGB6461*  *GGB4710 SGB6522*  *GGB6544 SGB9243*  *GGB6648 SGB9389*  *GGB9291 SGB14248*  *GGB9350 SGB14317*  *GGB9568 SGB14980*  *GGB9593 SGB15015*  *GGB9635 SGB15103*  *GGB9635 SGB15104*  *GGB9673 SGB15172*  *GGB9686 SGB15190*  *GGB9730 SGB15290*  *Hungatella hathewayi*  *Intestinimonas gabonensis*  *Lachnospiraceae bacterium 2 1 46FAA*  *Lachnospiraceae bacterium NSJ 46*  *Lachnospiraceae bacterium WCA 693 APC*  *MOT I*  *Lachnospiraceae unclassified SGB4924*  *Lactobacillus kalixensis*  *Leuconostoc pseudomesenteroides*  *Limosilactobacillus oris*  *Longicatena caecimuris*  *Methanosphaera stadtmanae*  *Microbacterium SGB53518*  *Monoglobus pectinilyticus*  *ParaPrevotella xylaniphila*  *Peptacetobacter hiranonis*  *Prevotella SGB1653*  *Prevotella SGB1680*  *Prevotella copri clade C*  *Prevotella copri clade D*  *Prevotella pectinovora*  *Prevotella sp P4 51*  *Prevotellamassilia timonensis*  *Rikenellaceae bacterium*  *Roseburia sp AM59 24XD*  *Ruminococcaceae unclassified SGB14835*  *Slackia piriformis*  *Streptococcus infantis* | - P221-PWY:   octane oxidation   - P562-PWY:   myo-inositol degradation I   - PWY-6318:   L-phenylalanine degradation IV  (mammalian. via side chain) |
| Endurance | *Acidaminococcus provencensis*  *Actinomyces SGB17154*  *Amedibacterium intestinale*  *Anaerococcus obesiensis*  *Anaerofustis stercorihominis*  *Anaerotruncus colihominis*  *Bacilli unclassified SGB6487*  *Blautia hydrogenotrophica*  *Blautia sp An81*  *Campylobacter hominis*  *Candidatus Heritagella gallinarum*  *Candidatus Neoruminococcus faecicola*  *Candidatus Pseudoruminococcus merdavium*  *Christensenella minuta*  *Clostridiales Family XIII Incertae Sedis*  *unclassified SGB3978*  *Clostridiales bacterium Marseille P5551*  *Coriobacteriia bacterium*  *Coriobacteriia unclassified SGB14764*  *Cryptobacterium curtum*  *Dielma fastidiosa*  *Erysipelotrichaceae bacterium 3 1 53*  *Ezakiella coagulans*  *Fenollaria timonensis*  *Frisingicoccus caecimuris*  *Fusobacterium mortiferum*  *GGB1109 SGB1423*  *GGB1215 SGB1581*  *GGB1407 SGB1931*  *GGB1455 SGB2018*  *GGB1456 SGB2019*  *GGB33665 SGB47358*  *GGB34228 SGB72916*  *GGB3583 SGB4799*  *GGB4277 SGB5832*  *GGB4569 SGB6312*  *GGB4584 SGB6338*  *GGB4667 SGB6456*  *GGB6580 SGB9301*  *GGB6608 SGB9342*  *GGB9610 SGB15045*  *GGB9618 SGB15064*  *GGB9646 SGB15123*  *GGB9778 SGB15398*  *Gordonibacter urolithinfaciens*  *Haemophilus paraphrohaemolyticus*  *Intestinimonas timonensis*  *Klebsiella oxytoca*  *Lachnoclostridium sp An118*  *Lachnoclostridium sp An138*  *Lachnospiraceae bacterium NSJ 29*  *Lactobacillus delbrueckii*  *Lactobacillus sp MRS 253 APC 2B*  *Leuconostoc carnosum*  *Leuconostoc lactis*  *Levyella massiliensis*  *Ligilactobacillus ruminis*  *Massilimaliae massiliensis*  *Massilimaliae timonensis*  *Megasphaera stantonii*  *Mesosutterella multiformis*  *Mitsuokella multacida*  *Mobiluncus SGB15488*  *Parabacteroides massiliensis*  *Pediococcus pentosaceus*  *Peptoniphilus senegalensis*  *Porphyromonas SGB1977*  *Porphyromonas bennonis*  *Porphyromonas somerae*  *Prevotella corporis*  *Prevotella sp 885*  *Saccharomyces cerevisiae*  *Schaalia turicensis*  *Streptococcus mitis*  *Streptococcus suis*  *Weissella cibaria* | - 12DICHLORETHDEG-PWY:   1.2-dichloroethane degradation   - 3-HYDROXYPHENYLACETATE-DEGRADATION-PWY:   4-hydroxyphenylacetate degradation   - PWY-5180:   toluene degradation I (aerobic) (via o-cresol)   - PWY-5392:   reductive TCA cycle II   - PWY-5415:   catechol degradation I (meta-cleavage pathway)   - PWY-5417:   catechol degradation III (orthocleavage pathway)   - PWY-5431:   aromatic compounds degradation   - PWY-5855:   ubiquinol-7 biosynthesis (early decarboxylation)   - PWY-6182:   superpathway of salicylate degradation   - PWY-6185:   4-methylcatechol degradation (ortho cleavage)   - PWY-6763:   salicortin biosynthesis   - PWY-6922:   L-N&delta;-acetylornithine biosynthesis   - PWY-7165:   L-ascorbate biosynthesis VIII  (engineered pathway)   - PWY-7245:   superpathway of NAD/NADP-NADH/NADPH interconversion (yeast)   - PWY-7268:   cytosolic NADPH production (yeast)   - PWY-7269:   mitochondrial NADPH production (yeast) |

**Supplementary Table 4.** Top correlations of species with fitness parameter values calculated by SparCC. Only correlations above 0.1 or top correlations per fitness parameter if none above 0.1 included.

| - **Fitness parameter** | - **Species** | - **r** |
| --- | --- | --- |
| - Fitness score | - *Phocaeicola vulgatus* | - 0.17 |
| - Fitness score | - *Bacteroides eggerthii* | - 0.11 |
| - Fitness score | - *Streptococcus thermophilus* | - 0.11 |
| - Fitness score | - *GGB3293 SGB4348* | - -0.19 |
| - Fitness score | - *Roseburia intestinalis* | - -0.17 |
| - VO_2_max | - *Bifidobacterium adolescentis* | - 0.18 |
| - VO_2_max | - *Bifidobacterium longum* | - 0.14 |
| - VO_2_max | - *Candidatus Cibiobacter qucibialis* | - 0.11 |
| - VO_2_max | - *Phocaeicola massiliensis* | - -0.13 |
| - VO_2_max | - *Lachnospira pectinoschiza* | - -0.13 |
| - VO_2_max | - *Eubacterium siraeum* | - -0.13 |
| - VO_2_max | - *Eubacterium siraeum* | - -0.13 |
| - Average power | - *Clostridiaceae bacterium* | - 0.09 |
| - Average power | - *Ruminococcus bromii* | - 0.08 |
| - Average power | - *Blautia wexlerae* | - 0.07 |
| - Average power | - *Clostridium innocuum* | - -0.08 |
| - Average power | - *GGB1227 SGB1600* | - -0.08 |
| - Average power | - *Roseburia inulinivorans* | - -0.07 |
| - Maximal power | - *Eubacterium rectale* | - 0.09 |
| - Maximal power | - *Blautia wexlerae* | - 0.09 |
| - Maximal power | - *GGB38171 SGB72433* | - 0.08 |
| - Maximal power | - *Phocaeicola massiliensis* | - -0.08 |
| - Maximal power | - *GGB9627 SGB15081* | - -0.07 |
| - Maximal power | - *Prevotella buccalis* | - -0.07 |

**Supplementary Table 5.** Correlations of functions with fitness parameter values calculated by SparCC. Only correlations above 0.1 or top correlations per fitness parameter if none above 0.1 included.

| - **Fitness parameter** | - **Function** | - **Species associated** | - **r** |
| --- | --- | --- | --- |
| - Fitness score | - PWY-7238: - sucrose biosynthesis II | - *Bifidobacterium adolescentis* | - 0.06 |
| - Fitness score | - PWY-1042: - glycolysis IV | - *Bacteroides vulgatus* | - -0.06 |
| - Fitness score | - PANTO-PWY: - phosphopantothenate biosynthesis I | - *Ruminococcus torques* | - -0.07 |
| - Fitness score | - PWY-5695: - inosine 5'-phosphate degradation | - *Fusicatenibacter saccharivorans* | - -0.07 |
| - VO_2_max | - PWY-6277: - superpathway of 5-aminoimidazole ribonucleotide biosynthesis | - *Blautia obeum* | - -0.06 |
| - VO_2_max | - PWY-8178: - pentose phosphate pathway (non-oxidative branch) II | - *Fusicatenibacter saccharivorans* | - 0.06 |
| - VO_2_max | - PWY-6317: - D-galactose degradation I (Leloir pathway | - *Faecalibacterium prausnitzii* | - 0.06 |
| - VO_2_max | - NONOXIPENT-PWY: - pentose phosphate pathway (non-oxidative branch) I | - *Fusicatenibacter saccharivorans* | - 0.06 |
| - VO_2_max | - PWY-7357: - thiamine phosphate formation from pyrithiamine and oxythiamine (yeast) | - *Blautia obeum* | - -0.06 |
| - VO_2_max | - PWY-5103: - L-isoleucine biosynthesis III | - *Prevotella copri* | - -0.07 |
| - VO_2_max | - PWY-7237: - myo-, chiro- and scyllo-inositol degradation | - *Ruminococcus torques* | - 0.07 |
| - VO_2_max | - PWY-6823: - molybdopterin biosynthesis | - *Blautia obeum* | - -0.07 |
| - VO_2_max | - PWY-6147: - 6-hydroxymethyl-dihydropterin diphosphate biosynthesis I | - *Prevotella copri* | - 0.07 |
| - VO_2_max | - LACTOSECAT-PWY: - lactose and galactose degradation I | - *Collinsella aerofaciens* | - -0.07 |
| - VO_2_max | - ARO-PWY: - chorismate biosynthesis I | - *Faecalibacterium prausnitzii* | - -0.09 |
| - VO_2_max | - PWY0-1296: - purine ribonucleosides degradation | - *Fusicatenibacter saccharivorans* | - -0.11 |
| - Average power | - PWY-6387: - UDP-N-acetylmuramoyl-pentapeptide biosynthesis I (meso-diaminopimelate containing) | - *Faecalibacterium prausnitzii* | - -0.06 |
| - Average power | - ARGSYN-PWY: - L-arginine biosynthesis I (via L-ornithine) | - *Blautia wexlerae* | - 0.07 |
| - Average power | - PWY-7238: - sucrose biosynthesis II | - *Blautia wexlerae* | - -0.08 |
| - Maximum power | - PWY66-429: fatty acid biosynthesis initiation (mitochondria) | - *Fusicatenibacter saccharivorans* | - -0.09 |
| - Maximum power | - PWY-6823: molybdopterin biosynthesis | - *Blautia obeum* | - -0.11 |

**Data availability statement**

The data supporting the findings of this study are openly available in the European Nucleotide Archive (ENA) at <https://www.ebi.ac.uk/ena/>, reference number PRJEB60692.
